# Symbiont switching and alternative resource acquisition strategies drive mutualism breakdown

**DOI:** 10.1101/242834

**Authors:** Gijsbert DA Werner, Johannes HC Cornelissen, William K Cornwell, Nadejda A Soudzilovskaia, Jens Kattge, Stuart A West, E Toby Kiers

## Abstract

Cooperative interactions among species, termed mutualisms, have played a crucial role in the evolution of life on Earth. However, despite key potential benefits to partners, there are many cases where two species cease to cooperate, and mutualisms break down. What factors drive the evolutionary breakdown of mutualism? We examined the pathways towards breakdowns of the mutualism between plants and arbuscular mycorrhizal (AM) fungi. Using a comparative approach, we identify ~25 independent cases of complete mutualism breakdown across global seed plants. We found that breakdown of cooperation was only stable when host plants either: (i) partner with other root symbionts or (ii) evolve alternative resource acquisition strategies. Our results suggest that key mutualistic services are only permanently lost if hosts evolve alternative symbioses or adaptations.

**Significance Statement:** Cooperative interactions among species – mutualisms – are major sources of evolutionary innovation. However, despite their importance, two species that formerly cooperated sometimes cease their partnership. Why do mutualisms breakdown? We asked this question in the partnership between arbuscular mycorrhizal (AM) fungi and their plant hosts, one of the most ancient mutualisms. We analyse two potential trajectories towards evolutionary breakdown of their cooperation, symbiont switching and mutualism abandonment. We find evidence that plants stop interacting with AM fungi when they switch to other microbial mutualists or when they evolve alternative strategies to extract nutrients from the environment. Our results show vital cooperative interactions can be lost - but only if successful alternatives evolve.

## Introduction

Mutualisms, cooperative partnerships among different species, have shaped much of Earth’s biodiversity, allowing organisms to outsource crucial functions like nutrition, cleaning, transport and defence (1, 2). Both theoretical and empirical work has provided us with a good understanding of the mechanisms, such as co-transmission and sanctions, that stabilise mutualism and maintain cooperation among species (3–5). Because of these mechanisms, beneficial interactions can be maintained over millions of years, and in some cases give rise to extreme mutualistic dependence (6, 7).

Despite reciprocal benefits, mutualisms do not always persist, and conflict among partners can remain. Theoretical and experimental work suggests that even when mutual benefits occur, fitness interests of both partners are generally not perfectly aligned, potentially selecting for cheaters and exploiters of mutualism (5, 8–11). This finding is further reinforced by the observation that in many mutualisms, there are mechanisms to evaluate partner quality and reward cooperation or sanction non-cooperative cheats (12–14). Furthermore, over ecological time, short-term breakdowns of cooperation in response to shifting environmental conditions, have been observed in many mutualisms, including plant rhizobial and mycorrhizal mutualisms, coral symbioses, protection and pollination mutualisms (15–19). Together, these observations raise the question in which conditions we should expect cooperation among species to fail, and partners in previously successful mutualisms to cease cooperating.

Even mutualisms that have become highly dependent over millions of years of co-evolution, have broken down in some occasions. This is the case, for example, when free-living fungi evolved from a previously lichenised lifestyle, or when parasitic moths evolved from pollinating ancestors (20–22). Yet, while we have a good understanding of why mutualistic cooperation is favoured, we lack a general understanding of the drivers of these evolutionary breakdowns of mutualisms. A number of non-exclusive reasons for the breakdown of mutualisms have been proposed (23, 24). Benefits provided by a mutualistic partner can become redundant through the evolution of alternative adaptations. In these cases, one of the partners switches from relying on another species to acquiring a function autonomously. For example, the evolution of large amounts of small-diameter pollen enabled the reversion back to an autonomous, wind-pollinated lifestyle in some angiosperms (25). A second trajectory occurs when one side of the interaction is replaced with a new mutualist species. While partner switching by definition leads to the evolution of a new partnership, the ancestral interaction is lost and thus a previously functional mutualism breaks down. This is illustrated in cases where plant species stop cooperating with birds and switch to insect pollination (26).

Our aim was to study the ancient and ubiquitous mutualism between plants and arbuscular mycorrhizal (AM) fungi to understand pathways towards mutualism breakdown. We focus on the plant-AM mutualism for three reasons. First, AM fungi (Glomeromycota) are among the most important terrestrial mutualists. AM fungi form extensive hyphal networks in the soil (up to 100 m cm^−3^ soil), providing plants with a key solution to the problem of extracting immobile nutrients, especially phosphorus (27). The partnership is crucial for plant growth, providing hosts with primarily phosphorus, but also nitrogen, water and trace elements (28). Second, even though the large majority of plants can be successfully colonised by AM fungi, 10-20% of plant species across a number of divergent clades do not interact with any AM fungi (27, 29). These repeated losses of the interaction, separated by millions of years of evolution, enable us to test general patterns and explanatory factors driving cooperation loss in a comparative framework. Third, the tools and databases allowing for broad comparative analyses are becoming available for plants, including a comprehensive phylogeny of seed plants (30), and large scale databases of plant traits including their association with AM fungi and other root symbionts (31–33).

In our analysis of the plant – AM mutualism, we take a plant-centric perspective. We are interested in cases where plants completely cease to interact with *all* AM fungi and where this lack of interaction persists over evolutionary time: *i.e*. where the loss of the interaction is not followed by host plant extinction. Thus, we do not study when plant-AM cooperation dissolves in the short-term due to ecological conditions, such as under high nutrient conditions. Rather, our aim is to first quantify stable losses of cooperation, and then test the importance of two types of evolutionary breakdown: partner switching and mutualism abandonment. By partner switching, we mean a situation where a host plant that ancestrally interacted with AM fungi, switched to interacting with a novel root symbiont with similar function and ceased interacting with AM fungi. We analyse switches to other mycorrhizal fungi, as well as to N_2_-fixing symbioses with rhizobial and *Frankia* bacteria (28, 34). We refer to mutualism abandonment, when plants have evolved an alternative strategy to acquire resource in a non-symbiotic way, for instance carnivory or cluster roots (35, 36).

## Results

### Evolutionary reconstruction of the plant-AM fungal mutualism

Our first aim was to quantify the evolutionary stability of the plant-AM mutualism, determining the number of losses of plant-AM interactions across the plant phylogeny. We compiled a global database of plant mycorrhizal fungal status across the seed plants (angiosperms and gymnosperms). We scored the reported interactions of plants with AM fungi in 3,736 plant species present in the most recent and comprehensive phylogeny of gymnosperms and angiosperms (30). We then established patterns of AM loss and gain using a Hidden Rate Model (HRM) approach to ancestral state reconstructions (37). This technique permits variation in the speed of binary character evolution so we can detect changes in rates of evolution, such as shifts in the evolutionary stability of plant-AM associations.

Our reconstructions revealed that the evolution of AM interactions across seed plants was best characterised by heterogeneity in speed of evolution: the best evolutionary model contains three different rate classes of evolution (Table 1). Specifically, we find strong evidence for the existence of an evolutionary class where AM interactions are strongly favoured (which we termed Stable AM), a class where an absence of AM interactions is strongly favoured (Stable Non-AM), and a class where AM interactions are evolutionarily labile (SI Figure 1).

**Table 1:** AICc-values and weights for all HRM models.

| Number of rate classes | Number of parameters | AICc | Δ-AICc | AICc-weight |
| --- | --- | --- | --- | --- |
| 1 | 2 | 3231.25 | 770.0 | 0.0% |
| 2 | 8 | 2597.78 | 136.5 | 0.0% |
| **3** | **14** | **2461.25** | **0** | **74.8%** |
| 4 | 20 | 2463.44 | 2.2 | 25.1% |
| 5 | 26 | 2473.79 | 12.5 | 0.1% |

*Table of the five different HRMs explored to analyse our AM fungal association data (see Methods). We used AICc-weights (corrected Akaike information criterion) to determine the model with the best fit (bold).*

Mapping these different evolutionary states back onto the phylogeny (SI Figure 2), we found that: (i) Association with AM fungi was likely the ancestral state of seed plants (99.6% likelihood); (ii) Stable AM fungal associations have been widely retained throughout the seed plants for over 350 million years, and represent the large majority of all historical and contemporary plant species and families (Table 2); (iii) some plant lineages evolve to either an evolutionarily labile state or a state where AM fungi are disfavoured (SI Figure 1, Table 2). Specifically, (iv) there have been an estimated ~25 evolutionary losses of the AM mutualism throughout the history of seed plants, found across 69 families (median over 100 bootstrap phylogenies 25.4, SD: 7.73). Which evolutionary trajectories are most important in explaining these breakdowns of cooperation among plants and AM fungi?

**Table 2:** Number of contemporary species and families in three AM Classes.

| Stability Class | Species numbers |  | Family numbers |  |
| --- | --- | --- | --- | --- |
|  | Best tree | Median | Best tree | Median |
| Stable AM | 2,616 | 2613 (SD 178) | 172 | 171 (SD 12) |
| Labile | 829 | 833 (SD 180) | 77 | 79 (SD 13) |
| Stable Non-AM | 291 | 288 (SD 21) | 8 | 7 (SD 3.7) |

*Numbers of contemporary species and families per AM stability class.*
*Median value and SD across 100 bootstrap phylogenies are indicated.*

### Symbiont switching and mutualism abandonment drive breakdown

We tested the hypotheses that AM loss is driven by shifts to other symbionts (partner switching) or by alternative adaptations for resource acquisition (abandonment). We generated a database of other major root symbionts with functional roles (providing phosphorus and nitrogen) similar to AM fungi. Specifically, based on a previously published database, we included presence or absence of a potential to interact with symbiotic N_2_-fixing bacteria (both rhizobial and *Frankia* bacteria) for all our host plant species (34, 38). We also included reported interactions with other mycorrhizal fungi (*i.e*. non-AM fungi that live in symbiotic association with plant roots). This included ectomycorrhizal (EM), ericoid mycorrhizal (ER), orchid mycorrhizal (ORM) and arbutoid mycorrhizal (ARB) fungi. All AM fungi belong to the division *Glomeromycota*, while other mycorrhizal fungi are only distantly related, belonging to a wide range of divisions, mainly *Basidiomycota* (ECM, ARB and ORM, some ER) and *Ascomycota* (some ECM and ORM and most ER). Some plant species interact with multiple types of mycorrhizal fungi (28, 39).

We scored our species for the reported presence of alternative resource acquisition strategies. These included parasitism as a plant strategy (both plants parasitising other plants and full mycoheterotrophs, *i.e*. plants parasitising mycorrhizal fungi) (40, 41), carnivory (35) and cluster roots (36) (Figure 1). These strategies have in common that they represent alternative solutions to the problem of acquiring scarce mineral resources: they acquire resources by seizing them from other organisms (plant parasitism), through direct predation (carnivorous plants), or through investing in a unique root architecture characterised by a high density of finely-branched roots and root hairs, known as cluster roots (Figure 1). To study congruence between losses of AM interactions and alternative strategies, we again performed ancestral state reconstructions, to study the origins of: (i) other symbionts (*i.e*. non-AM mycorrhizal fungal symbionts or symbiotic N_2_-fixation), which were present in 820 of our 3,736 plant species; and (ii) alternative resource acquisition strategies, present in 109 plant species.

**Figure 1:**
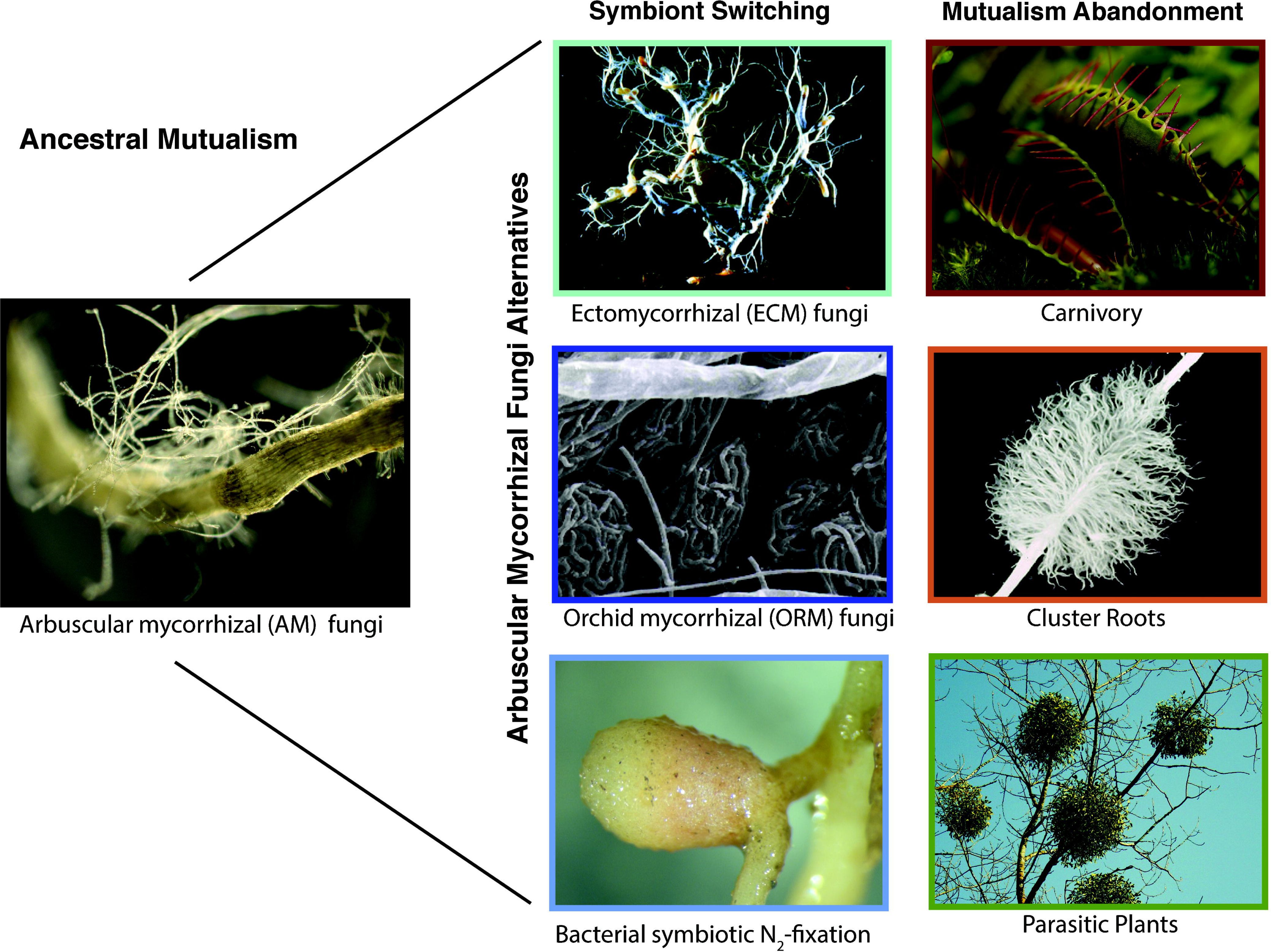
We explored the evolution of various alternatives to interacting with AM fungi, the ancestral state of seed plants. Examples of six important alternatives we considered are depicted, with columns indicating two potential pathways towards evolutionary breakdown of the plant-AM mutualism. In both pathways, the ancestral mutualism with AM fungi breaks down. Coloured borders match coloured bars in Figure 2, indicating distribution of these traits across global seed plants.

We found a high degree of congruence between the different origins of AM losses and of various AM-alternatives (SI Table 1, SI Figures 3-9). To study this quantitatively, we compared models of dependent *vs*. independent evolution (42), analysing the relationship between AM loss and presence of alternative partners or alternative resources acquisition strategies. We studied a binary variable coding for presence of any AM alternative and found that a dependent model of evolution vastly outperformed an independent model (Δ-AICc 428.90, AICc-weight 99.9%). This means that over evolutionary time, AM loss (shift from the bottom left plane to the top right in the transitions matrix, Figure 2) is strongly associated with the presence of another mycorrhizal fungal partner, or alternative resource strategy. Thus, partner switching and mutualism abandonment are important in enabling evolutionary breakdown of the ancestral plant-AM fungal mutualism throughout the seed plants.

**Figure 2:**
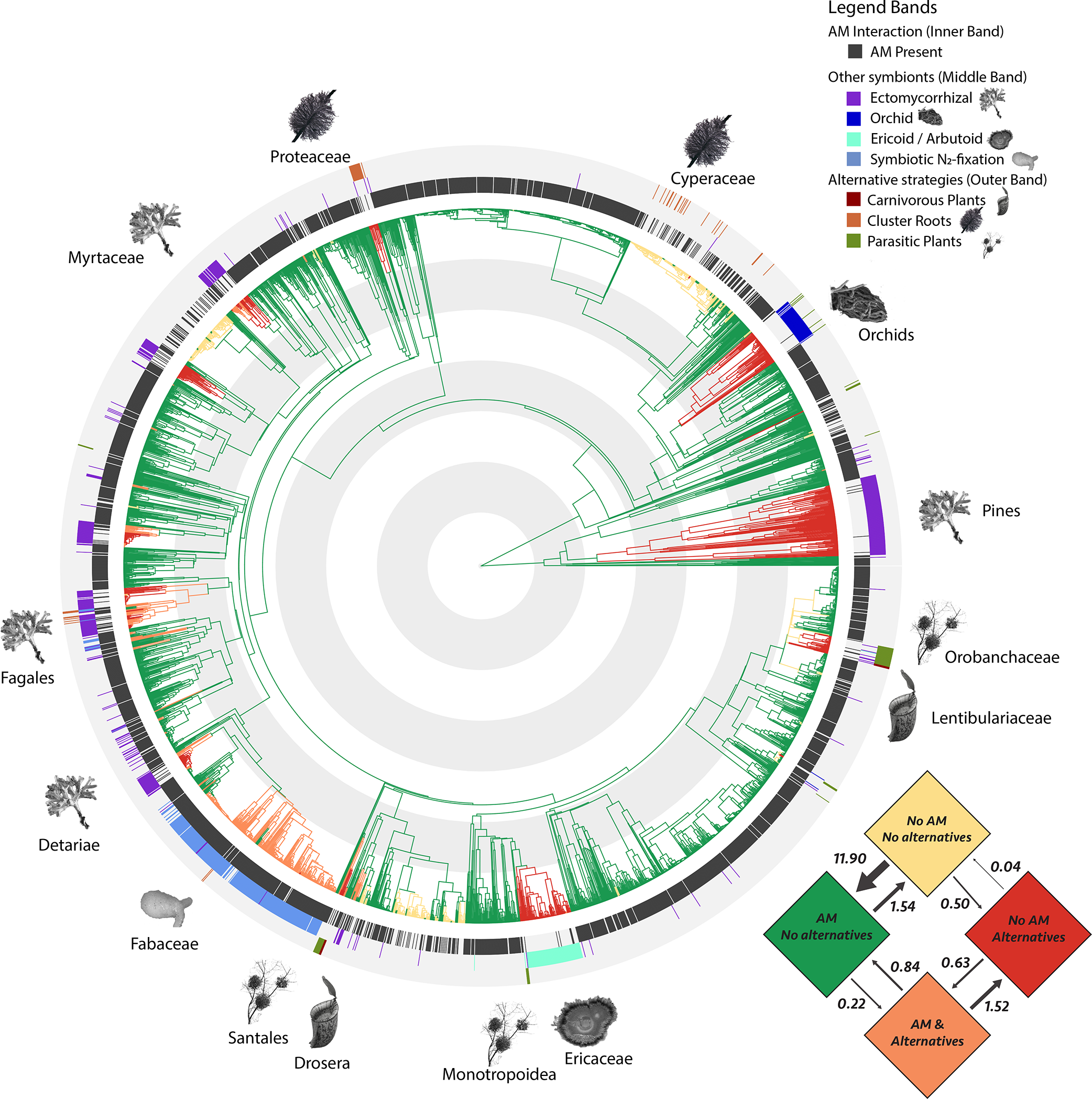
Transition rates and ancestral state reconstruction of the dependent evolutionary model for plant AM status and AM-alternatives. The four potential evolutionary states are represented by colours in the transition matrix and on the seed plant phylogeny. Transition rates are expressed as number of transitions per 100 million years per lineage. The ancestral state is AM presence with no AM-alternatives (green), red indicates the switch to one of the AM alternatives (i.e. another mycorrhizal fungus, symbiotic N_2_-fixation, parasitism, carnivory or cluster roots). From inside to outside, coloured bands around the phylogeny indicate the presence or absence of (i) AM interactions, (ii) other root symbionts and (iii) alternative resource acquisition strategies. Key clades that have lost AM fungal interactions are indicated with schematic images of their evolved alternatives. Grey and white concentric circles indicate 50 million years. An expanded version with fully legible species labels for all 3,736 species is available online.

More specifically, from the inferred transition matrix and associated ancestral state reconstruction (Figure 2), we conclude that: (i) The AM mutualism is generally highly stable - transition rates towards the AM state (green) are about ten times as high as losses (from green to yellow) (ii) AM fungal loss is only stable when an alternative is present (orange to red) and not without (green to yellow). (iii) While evolutionary stability is high when plants associate with either AM fungal symbionts (green state) or an alternative symbiont or acquisition strategy (red), having neither (yellow) is evolutionarily unstable. For instance, all the origins of this type (*e.g*. in the Brassicales) have occurred relatively recently in evolutionary terms (within the last 30 million years). (iv) Similarly, it is evolutionarily less stable to have both AM and an alternative simultaneously (orange).

Our reconstructions show that both the evolutionary scenario of initial AM loss followed by alternative strategy evolution and the reverse order are possible. Initial acquisition of an AM-alternative (move from green to orange state), in some cases may have resulted in released selection to maintain the AM interaction, allowing for its subsequent evolutionary breakdown (orange to red). In other cases, the AM interaction was lost first (yellow state), for instance through symbiosis gene loss (43), and survival of host plants was subsequently favoured when rapidly evolving an AM alternative state. Thus, our analysis indicates there is no single dominant trajectory in the transition from an AM plant to a stable non-AM plant, but that both routes can occur.

### Sensitivity Analyses

To verify the robustness of our results, we considered the sensitivity of our main conclusions to two forms of uncertainty, (1) phylogenetic uncertainty and (2) uncertainty in the underlying AM data. We analysed phylogenetic uncertainty by replicating our initial AM fungal reconstruction analysis over 100 bootstrap phylogenies, and found highly similar relative loss rates of the plant AM-interaction throughout 100% of our bootstrap replicates (SI Figure 10) and highly similar ancestral state reconstructions (SI Figure 11). We also found that across the 100 bootstrap phylogenies a dependent model of evolution always outperformed an independent model (mean Δ-AICc 390.52). This further confirms the deep evolutionary link between AM loss and the evolution of other symbionts and resource acquisition strategies regardless of the details of the phylogenies used (SI Figure 12).

A second main source of uncertainty is in the AM status of plants. This is because AM fungi are notoriously difficult to score: it is easy to misidentify other fungi as AM fungi (false positive) or to miss AM hyphae (false negative). To address this, we implemented a re-simulation approach which takes into account the number of independent reports of AM status, and allows us to test separate false positive and false negative rates for these underlying reports. We found that even if one in four of the AM reports in our database is incorrect (*e.g*. a saprotrophic fungus) while simultaneously 25% of our AM absence reports in fact were mycorrhizal, we still draw highly similar conclusions (SI Figure 13, SI Table 2). Therefore, even if we assume the underlying mycorrhizal data are of very poor quality, we recover qualitatively highly similar patterns. Thus, overall, we conclude that all our main conclusions are robust to substantial phylogenetic and data uncertainty.

As a final analysis, we compared our results with a recent comparative analysis of plant-mycorrhizal symbioses (29). This analysis used an alternative scoring approach that divided plant species in four categories: AM plants, non-mycorrhizal plants (NM-plants), ECM plants, and plants that are commonly found in either AM or NM states (AMNM-plants), and found that transitions from AM towards NM states primarily go through the AMNM state. We re-confirmed this result, in that we find in our best HRM-model of plant-AM interactions that plants transition through the labile state to the stable non-AM state, where the loss of plant-AM mutualism becomes evolutionarily entrenched (SI Figure 1). We also find that the species-level percentage of observations with AM-presence has a median value of 100% (SE: 0.89%; mean 83.4%) for species inferred to be in the stable AM class and 0% in the stable non-AM class (SE: 0.73%; mean 1.58%), while in the evolutionarily labile class this is 16.7% (SE: 1.57%; mean 22.0%; SI Figure 14). This indicates that the labile state inferred under our deep evolutionary model effectively recovers the notion of an AMNM presupposed by Maherali *et al*. While their analysis allows for direct inclusion of AMNM and ECM states, with our approach of binary coding the presence or absence of AM and other mycorrhizal interactions we can answer different questions: (i) It allows us to infer the *variation* in loss rate of the AM mutualism across seed plant evolutionary history (which is only possible in the HRM-framework for binary traits (37)) (ii) Rather than *a priori* defining an intermediate state, it allows us to verify if an evolutionarily labile state is actually inferred in our best model. (iii) It allows us to study the dependent evolution of AM and other mycorrhizal interactions as separate traits. This is especially important because, while rare, dual colonisation of plants by two types simultaneously is possible and could represent an important evolutionary intermediary state, as confirmed by our analysis (Figure 2). (4) It allows us to include in our analysis not just ECM fungi, but also other root symbionts such as symbiotic N_2_-fixation, ericoid (ERM) and orchid (ORM) mycorrhizal fungi, which turned out to be drivers of major evolutionary losses of the plant-AM mutualism (Figure 2).

## Discussion

Our analyses revealed that the ancient and ubiquitous plant–AM fungal mutualism has broken down in ~25 cases across the seed plants. We found that stable and persistent mutualism breakdown is driven both by acquisitions of other root symbionts (partner switching) and by the evolution of alternative non-symbiotic resource acquisition strategies (mutualism abandonment).

These results in turn raise the question of what underlying ecological factors favour transitions to these alternative solutions, and the mechanisms that enable them. Mechanistically, an important step is likely the loss of key genes in the ‘symbiotic toolkit’ encoding crucial root mutualism effectors (43, 44). This must either be followed or preceded by molecular evolution in the genes encoding alternative symbioses or resource acquisition traits. Ecologically, these alternatives can potentially be favoured by a range of ultimate factors, such as environmental change, habitat shifts (for instance to high-nutrient soils), migration, invasion or partner abundance (22, 24, 45–49), although discriminating these over deep evolutionary time is challenging. One hypothesis is that switching from the AM nutrient uptake strategy to rarer alternative strategies has enabled plants to compete in a range of (micro)habitats. Evolution of carnivory in temperate swamps (35), cluster-roots in extremely phosphorus-impoverished soils (36), cold-resistant ectomycorrhizal interactions in lower temperature habitats (50) and ericoid mycorrhizal fungi in resource-poor heath lands (51) has helped host plants to thrive in environments where the more common AM interaction is a less successful solution to obtain nutrients.

We emphasise that our estimate of ~25 breakdowns represents a conservative lower bound, since we study plants that stopped interacting with *all* AM fungal species and subsequently persisted over evolutionary times. The number of breakdowns of plant mutualism with specific AM fungal species or lineages while cooperation with other AM continued, is likely to be considerably higher. More generally, studies that analyse breakdowns of symbiotic interactions with entire taxa of organisms - such as among corals and any photosynthetic dinoflagellates (52, 53) – will underestimate the number of breakdowns with specific symbiont species.

We find that dual symbioses – simultaneously being able to interact with two types of root symbionts - is unlikely to be evolutionarily stable (Figure 2). This is a different pattern compared to what is documented in insect endosymbioses, which often acquire secondary partners while retaining the ancestral mutualism (54, 55). In insects, maintenance of two endosymbionts could be favoured by different microbial partners subsequently specialising on different mutualistic functions (56, 57). In root symbioses, nutritional benefits provided by AM fungi and by alternative root symbionts may often be too similar to outweigh the costs of maintaining them both. For instance, while AM fungi are thought to provide primarily phosphorus, they also contribute nitrogen to their hosts (58). This could help explain why plants only rarely associate with both ECM fungi and AM fungi simultaneously (131 species in our dataset). However, our reconstructions suggest that such dual symbioses can be a transitory state on the path towards a complete switch and breakdown of the original mutualism (Figure 2), as was previously hypothesised (59). Other root symbionts may provide more complementary benefits to their plant hosts, which could select for maintenance of dual symbioses. For instance, AM fungi and N_2_-fixing rhizobial bacteria are often thought to provide complementary benefits to their legume hosts (60), although a meta-analysis did not generally find synergistic effects on host growth (61).

If breakdown of the AM fungal mutualism is driven by acquisition of other root symbionts or alternative resources strategies, how can we explain plants that have neither AM fungi nor an alternative (yellow in Figure 2)? Recently, a member of the Brassicaceae, a family generally lacking mycorrhizal symbionts, was found to engage in a specific and beneficial interaction with fungi from the order Helotiales (Ascomycota), which provides soil nutrients (phosphate) to their hosts (62). While we do not know how widespread this phenomenon yet is, it raises the intriguing possibility that some of our species without AM fungi have in fact evolved interactions with yet unknown beneficial root symbionts functionally similar to mycorrhizal fungi. This would further strengthen the relationship we observed among AM loss and switches to alternative symbionts. Another, non-mutually exclusive possibility is that plants abandoning the AM fungal mutualism - without evolving alternatives - are likely to go extinct after an evolutionarily short period of time, or rapidly re-establish the mutualism. In line with this, all cases of AM breakdown not coupled to an alternative (yellow state; Figure 2), have evolved fairly recently (<30 MYA), compared to many much older losses associated with symbiont switching or alternative strategies (e.g. the switch to ECM fungi in Pines, more than 200 MYA).

An alternative potential reason for mutualism breakdown is when cheaters, low quality partners or parasites, arise in one of the partner lineages (23, 24). This can drive the interaction from mutual benefit to parasitism, and cause the other partner to abandon the interaction (9). Theory and empirical work suggests that hosts are particularly vulnerable to cheating when partners are acquired directly from the environment, like AM fungi (4, 5, 63–65). However, in bacteria, phylogenetic work has shown that while transitions towards cooperative states are common, loss of mutualist status is rare for bacterial symbionts (66, 67). When these losses occur, bacteria are more likely to revert to a free-living state than to become parasites (66, 67). In our case, such a reversion to a free-living state would correspond to a plant evolving an abiotic adaptation to replace AM fungi, such as cluster roots. While most of our ~25 losses can be explained in terms of symbiont switches or alternative resource strategies (Figure 2), some of the switches to other root symbionts or resource strategies we observed could initially have been driven by the fitness cost of parasitic AM fungi.

Our analyses show that cooperation among plants and AM fungi has generally persisted in a highly stable state for over 350 million years. This illustrates the importance of mutualistic services provided by AM fungi for most host plant species. Yet, even ancient and versatile mutualists like AM fungi can be completely and permanently lost in the right circumstances: we estimate this happened ~25 times. In general, mutualistic partnerships allow organisms to outsource crucial functions to other species, thereby obtaining these services more efficiently (5). Our results highlight how a key mutualistic service like nutrient acquisition is only permanently lost if hosts evolve either symbiotic or abiotic alternatives to obtain these functions.

## Methods

More detailed Extended Methods can be found in the online Supporting Information.

### Mycorrhizal status database

We compiled our database of reported plant mycorrhizal status by obtaining data from both primary literature and publicly accessible databases. Our full data source list, as well as our scoring criteria can be found in the Extended Methods (Supporting Information). Our analysed database contained data for a total of 3,736 spermatophyte species (3,530 angiosperms, 206 gymnosperms, 61 orders, 230 families and 1,629 genera) that overlapped with the phylogeny used in our analysis (30), is available online (Supporting Data 1).

### Reconstruction of the evolution of AM interactions

We used a Hidden Markov Model approach called ‘Hidden Rate Models’ (HRMs) which allows for heterogeneity in the loss and gain rates of a binary trait across a phylogeny (37). We used the R-package *corHMM* (37)(version 1.18) in R 3.2.3 to analyse our mycorrhizal data and explored HRMs with one to five rate classes, using AICc-weights to select the best HRM among this family of candidate models (Table 1). We used the marginal method to perform ancestral state reconstructions and employed Yang’s method to compute the root state (68). We *a posteriori* labelled the three rate classes under the best model ‘Stable AM’, ‘Labile’ and ‘Stable Non-AM’ (SI Figure 1).

### Database Alternative Resource Acquisition Strategies

We generated a second database for all our 3,376 analysed species and scored each species for the presence or absence of three main resource strategies, which each represent an alternative way of extracting minerals from the environment: carnivory (35), parasitism (40, 41) and cluster roots (36). Based on our previously generated database of plant species associating with symbiotic nitrogen-fixing bacteria (34, 38), we also assigned all analysed species a binary symbiotic nitrogen-fixation status. We describe our full data sources and scoring procedures in the Extended Methods.

### Correlated evolution of AM interactions and AM-alternatives

We generated HRM-models (37) of both non-AM mycorrhizal fungi and adaptations for resource acquisition (SI Table 1), plotted them onto our AM ancestral state reconstruction and visually identified the origins of these AM-alternatives (SI Figure 3-9). We then tested the potential for correlated evolution among AM fungi, other mycorrhizal fungi and resource acquisition adaptations. Using AIC-criteria, we compared models of dependent and independent evolution (42, 69) among the binary variables AM and AM-alternatives. We utilised the Maximum Likelihood implementation of the Discrete-module in *BayesTraits V2*, and constrained the ancestral node of the phylogeny to have AM fungi but none of the alternatives, as that is what our previous analyses had revealed (SI Figures 3-9).

### Sensitivity analysis to phylogenetic and data uncertainty

We studied the robustness of our main conclusions to two main sources of uncertainty: phylogenetic uncertainty and uncertainty in the underlying mycorrhizal data. We reran our key models (three rate class HRM and correlated evolution models in *BayesTraits*) across hundred bootstrap phylogenies (30) (SI Figure 11 and 12). To test for effects of data uncertainty, we used a resimulation approach that for each species takes into the number of observations of a given mycorrhizal state and simulates different error rates for underlying mycorrhizal observations (SI Figure 13, SI Table 2). We detail our full approach to the sensitivity analyses in the Extended Methods.

### Data Availability

Our full dataset, including number of reports of various mycorrhizal states across databases (see Extended Methods), our resulting assignment of AM, ECM, ORM, ER, ARB and symbiotic N_2_-fixation states, and our assignment of alternative resources acquisition strategies (carnivory, parasitism, mycoheterotrophy) is available online (Supporting Data 1).

## Acknowledgements

We thank SURFsara (www.surf-sara.nl) for support in using the Lisa Computing Cluster. The AM and AM-alternative illustrations in Figures 1 and 2 are based on figures in the public domain (CC0) with the exception of the illustrations for ericoid (courtesy Dr. David Midgley, CC-BY-SA) and arbuscular mycorrhizae (courtesy dr. Yoshihiro Kobae), root nodules (courtesy Dr. Euan James) and the illustrations for cluster roots (70) and orchid mycorrhizae (71) which were reprinted with permission from the respective publishers. The study has been supported by the TRY initiative on plant traits (http://www.trydb.org). GDAW was funded by a Royal Society Newton International Fellowship and a Junior Research Fellowship at Balliol College Oxford. ETK was funded by Netherlands Organisation for Scientific Research Grants 836.10.001 and 864.10.005 and European Research Council ERC Grant Agreement 335542.

## References

1. Bronstein JL (2015) Mutualism (Oxford University Press, Oxford, UK).

2. Weber MG, Agrawal A a. (2014) Defense mutualisms enhance plant diversification. Proc Natl Acad Sci 111(46):16442–16447.

3. West SA, Griffin AS, Gardner A (2007) Evolutionary Explanations for Cooperation. Curr Biol 17(16):R661–R672.

4. Sachs JL, Mueller UG, Wilcox TP, Bull JJ (2004) The evolution of cooperation. Q Rev Biol 79(2):135–160.

5. Leigh EG (2010) The evolution of mutualism. J Evol Biol 23(12):2507–2528.

6. Bennett GM, Moran NA (2015) Heritable symbiosis: The advantages and perils of an evolutionary rabbit hole. Proc Natl Acad Sci 112(33):10169–10176.

7. Neiman M, Lively CM, Meirmans S (2017) Why Sex? A Pluralist Approach Revisited. Trends Ecol Evol. doi:10.1016/j.tree.2017.05.004.

8. Herre EA, West SA (1997) Conflict of interest in a mutualism: documenting the elusive fig wasp-seed trade-off. Proc R Soc London Ser B, Biol Sci 264:1501–1507.

9. Ghoul M, Griffin AS, West S a (2014) TOWARD AN EVOLUTIONARY DEFINITION OF CHEATING. Evolution (N Y) 68(2):318–331.

10. Bao, Addicott (1998) Cheating in mutualism: defection of Yucca baccata against its yucca moths. Ecol Lett 1(3):155–159.

11. Sachs JL, Ehinger MO, Simms EL (2010) Origins of cheating and loss of symbiosis in wild Bradyrhizobium. J Evol Biol 23(5):1075–1089.

12. Kiers ET, et al. (2011) Reciprocal Rewards Stabilize Cooperation in the Mycorrhizal Symbiosis. Science (80- ) 333(6044):880–882.

13. Kiers ET, Rousseau RA, West SA, Denison RF (2003) Host sanctions and the legume–rhizobium mutualism. Nature 425(6953):78–81.

14. Wang R-W, Dunn DW, Sun BF (2014) Discriminative host sanctions in a fig–wasp mutualism. Ecology 95(5):1384–1393.

15. Husband R, Herre EA, Young JPW (2002) Temporal variation in the arbuscular mycorrhizal communities colonising seedlings in a tropical forest. FEMS Microbiol Ecol 42(1):131–6.

16. Baker DM, Freeman CJ, Wong JCY, Fogel ML, Knowlton N (2018) Climate change promotes parasitism in a coral symbiosis. ISME J. doi:10.1038/s41396-018-0046-8.

17. Batterman SA, et al. (2013) Key role of symbiotic dinitrogen fixation in tropical forest secondary succession. Nature 502(7470):224–227.

18. Ohm JR, Miller TEX (2014) Balancing anti-herbivore benefits and anti-pollinator costs of defensive mutualists. Ecology 95(10):2924–2935.

19. Menzel F, Kriesell H, Witte V (2014) Parabiotic ants: the costs and benefits of symbiosis. Ecol Entomol 39(4):436–444.

20. Lutzoni F, Pagel M, Reeb V (2001) Major fungal lineages are derived from lichen symbiotic ancestors. Nature 411(6840):937–940.

21. Pellmyr O, Leebens‐Mack J (2000) Reversal of Mutualism as a Mechanism for Adaptive Radiation in Yucca Moths. Am Nat 156(S4):S62–S76.

22. Kawakita A, Mochizuki K, Kato M (2015) Reversal of mutualism in a leafflower-leafflower moth association: The possible driving role of a third-party partner. Biol J Linn Soc 116(3):507–518.

23. Sachs J, Simms E (2006) Pathways to mutualism breakdown. Trends Ecol Evol 21(10):585–592.

24. Kiers ET, Palmer TM, Ives AR, Bruno JF, Bronstein JL (2010) Mutualisms in a changing world: an evolutionary perspective. Ecol Lett 13(12):1459–1474.

25. Culley TM, Weller SG, Sakai AK (2002) The evolution of wind pollination in angiosperms. Trends Ecol Evol 17(8):361–369.

26. Whittall JB, Hodges S a (2007) Pollinator shifts drive increasingly long nectar spurs in columbine flowers. Nature 447(7145):706–709.

27. Parniske M (2008) Arbuscular mycorrhiza: the mother of plant root endosymbioses. Nat Rev Microbiol 6(10):763–775.

28. van der Heijden MGA, Martin FM, Selosse M-A, Sanders IR (2015) Mycorrhizal ecology and evolution: the past, the present, and the future. New Phytol 205(4):1406–1423.

29. Maherali H, Oberle B, Stevens PF, Cornwell WK, McGlinn DJ (2016) Mutualism Persistence and Abandonment during the Evolution of the Mycorrhizal Symbiosis. Am Nat 188(5):E113–E125.

30. Zanne AE, et al. (2014) Three keys to the radiation of angiosperms into freezing environments. Nature 506(7486):89–92.

31. Kattge J, et al. (2011) TRY - a global database of plant traits. Glob Chang Biol 17(9):2905–2935.

32. Wang B, Qiu Y-L (2006) Phylogenetic distribution and evolution of mycorrhizas in land plants. Mycorrhiza 16(5):299–363.

33. Akhmetzhanova AA, et al. (2012) A rediscovered treasure: mycorrhizal intensity database for 3000 vascular plant species across the former Soviet Union. Ecology 93(3):689–690.

34. Werner GDA, Cornwell WK, Sprent JI, Kattge J, Kiers ET (2014) A single evolutionary innovation drives the deep evolution of symbiotic N2-fixation in angiosperms. Nat Commun 5:4087.

35. Ellison AM, Gotelli NJ (2001) Evolutionary ecology of carnivorous plants. Trends Ecol Evol 16(11):623–629.

36. Neumann G, Martinoia E (2002) Cluster roots – an underground adaptation for survival in extreme environments. Trends Plant Sci 7(4):162–167.

37. Beaulieu JM, O’Meara BC, Donoghue MJ (2013) Identifying hidden rate changes in the evolution of a binary morphological character: the evolution of plant habit in campanulid angiosperms. Syst Biol 62(5):725–37.

38. Werner GDA, Cornwell WK, Cornelissen JHC, Kiers ET (2015) Evolutionary signals of symbiotic persistence in the legume–rhizobia mutualism. Proc Natl Acad Sci 112(33):10262–10269.

39. Martin F, Kohler A, Murat C, Veneault-Fourrey C, Hibbett DS (2016) Unearthing the roots of ectomycorrhizal symbioses. Nat Rev Microbiol. doi:10.1038/nrmicro.2016.149.

40. Merckx VS (2013) Mycoheterotrophy ed Merckx V (Springer New York, New York, NY) doi:10.1007/978-1-4614-5209-6.

41. Těšitel J (2016) Functional biology of parasitic plants : a review. Plant Ecol Evol 149(1):5–20.

42. Pagel M (1994) Detecting Correlated Evolution on Phylogenies: A General Method for the Comparative Analysis of Discrete Characters. Proc R Soc B Biol Sci 255(1342):37–45.

43. Delaux P-M, et al. (2014) Comparative Phylogenomics Uncovers the Impact of Symbiotic Associations on Host Genome Evolution. PLoS Genet 10(7):e1004487.

44. Bravo A, York T, Pumplin N, Mueller LA, Harrison MJ (2016) Genes conserved for arbuscular mycorrhizal symbiosis identified through phylogenomics. Nat Plants 2(2):15208.

45. Oliver KM, Degnan PH, Burke GR, Moran N a (2010) Facultative symbionts in aphids and the horizontal transfer of ecologically important traits. Annu Rev Entomol 55:247–266.

46. Weese DJ, Heath KD, Dentinger BTM, Lau J a. (2015) Long-term nitrogen addition causes the evolution of less-cooperative mutualists. Evolution (N Y) 69(3):631–642.

47. Gutiérrez-Valencia J, Chomicki G, Renner SS (2017) Recurrent breakdowns of mutualisms with ants in the neotropical ant-plant genus Cecropia (Urticaceae). Mol Phylogenet Evol 111:196–205.

48. Chomicki G, Renner SS (2017) Partner abundance controls mutualism stability and the pace of morphological change over geologic time. Proc Natl Acad Sci 114(15):3951–3956.

49. Sheldrake M, et al. (2017) A phosphorus threshold for mycoheterotrophic plants in tropical forests. Proc R Soc B Biol Sci 284(1848):20162093.

50. Treseder KK, et al. (2014) Evolutionary histories of soil fungi are reflected in their large-scale biogeography. Ecol Lett 17(9):1086–1093.

51. Read D (1996) The Structure and Function of the Ericoid Mycorrhizal Root. Ann Bot 77(4):365–374.

52. Kitahara M V., Cairns SD, Stolarski J, Blair D, Miller DJ (2010) A comprehensive phylogenetic analysis of the scleractinia (Cnidaria, Anthozoa) based on mitochondrial CO1 sequence data. PLoS One 5(7). doi:10.1371/journal.pone.0011490.

53. Barbeitos MS, Romano SL, Lasker HR (2010) Repeated loss of coloniality and symbiosis in scleractinian corals. Proc Natl Acad Sci 107(26):11877–11882.

54. Hudson LN, et al. (2016) The database of the PREDICTS (Projecting Responses of Ecological Diversity In Changing Terrestrial Systems) project. Ecol Evol (September 2016):145–188.

55. Bennett GM, McCutcheon JP, MacDonald BR, Romanovicz D, Moran NA (2014) Differential genome evolution between companion symbionts in an insect-bacterial symbiosis. MBio 5(5):e01697–14.

56. Douglas AE (2016) How multi-partner endosymbioses function. Nat Rev Microbiol 14(12):731–743.

57. Szabó G, et al. (2017) Convergent patterns in the evolution of mealybug symbioses involving different intrabacterial symbionts. ISME J 11(3):715–726.

58. Govindarajulu M, et al. (2005) Nitrogen transfer in the arbuscular mycorrhizal symbiosis. Nature 435(7043):819–

59. Brundrett MC, Tedersoo L (2018) Evolutionary history of mycorrhizal symbioses and global host plant diversity. New Phytol. doi:10.1111/nph.14976.

60. Larimer AL, Clay K, Bever JD (2014) Synergism and context dependency of interactions between arbuscular mycorrhizal fungi and rhizobia with a prairie legume. Ecology 95(4):1045–1054.

61. Larimer AL, Bever JD, Clay K (2010) The interactive effects of plant microbial symbionts: a review and meta-analysis. Symbiosis 51(2):139–148.

62. Almario J, et al. (2017) Root-associated fungal microbiota of nonmycorrhizal Arabis alpina and its contribution to plant phosphorus nutrition. Proc Natl Acad Sci 114(44):E9403–E9412.

63. Porter SS, Simms EL (2014) Selection for cheating across disparate environments in the legume-rhizobium mutualism. Ecol Lett 17(9):1121–1129.

64. Salem H, Onchuru TO, Bauer E, Kaltenpoth M (2015) Symbiont transmission entails the risk of parasite infection. Biol Lett 11(12):20150840.

65. Gordon BR, et al. (2016) Decoupled genomic elements and the evolution of partner quality in nitrogen-fixing rhizobia. Ecol Evol 6(5):1317–1327.

66. Sachs JL, Skophammer RG, Regus JU (2011) Evolutionary transitions in bacterial symbiosis. Proc Natl Acad Sci 108(Supplement_2):10800–10807.

67. Sachs JL, Skophammer RG, Bansal N, Stajich JE (2013) Evolutionary origins and diversification of proteobacterial mutualists. Proc R Soc B Biol Sci 281(1775):20132146–20132146.

68. Yang Z (2006) Computational Molecular Evolution (Oxford University Press, Oxford, UK).

69. Pagel M (1999) Inferring the historical patterns of biological evolution. Nature 401(October 1999):877–884.

70. Armstrong A (2015) Ecophysiology: Root-cluster control. Nat Plants 1(2):15009.

71. Beyrle HF, Smith SE, Franco CMM, Peterson RL (1995) Colonization of Orchis morio protocorms by a mycorrhizal fungus: effects of nitrogen nutrition and glyphosate in modifying the responses. Can J Bot 73(8):1128–1140.

